# Population Pharmacokinetics of an Anti-PD-1 Antibody Camrelizumab in Patients with Multiple tumor types and model informed dosing strategy

**DOI:** 10.1101/2020.06.30.180117

**Authors:** Chen-yu Wang, Chang-cheng Sheng, Guang-li Ma, Da Xu, Xiao-qin Liu, Yu-ya Wang, Li Zhang, Chuan-liang Cui, Bing-he Xu, Yu-qin Song, Jun Zhu, Zheng Jiao

## Abstract

**Objective:** Camrelizumab, a programmed cell death 1 (PD-1) inhibitor, has been approved for the treatment of relapsed or refractory classical Hodgkin lymphoma. The aim of this study was to perform a population pharmacokinetics (PK) analysis of camrelizumab to quantify the impact of patient characteristics on PK and to investigate the appropriateness of flat dose in the dosing regimen.

**Methods:** A total of 3298 camrelizumab concentrations from 133 patients from four studies were analyzed using nonlinear mixed effects modeling. Covariate model building was conducted using stepwise forward addition and backward elimination. Monte Carlo simulation was conducted to compare exposures of 200 mg and 3 mg/kg every 2-week regimens.

**Results:** The PK of camrelizumab were adequately described by a two-compartment model with parallel linear and nonlinear clearances. Baseline albumin had significant effects on linear clearance, and weight had effects on inter-compartmental clearance. Moreover, 200 mg and 3 mg/kg regimens provide similar exposure distributions with no advantage to either dosing approach.

**Conclusion:** Population PK analysis provided an integrated evaluation of the impact of albumin and weight on the PK of camrelizumab. It also provided evidence that neither the flat-dose nor the weight-based dose regimen was advantageous over the other for most patients with tumors.

## 1 Introduction

The programmed cell death 1 (PD-1) pathway plays a critical role in maintaining an immunosuppressive tumor microenvironment, and blockade of the PD-1 pathway has become the key component of cancer immunotherapy.^1^ Camrelizumab (SHR-1210, AiRuiKa™) is a humanized high-affinity IgG4-kappa monoclonal antibody (mAb) to PD-1.^2^ In May 2019, the National Medical Products Administration of China approved camrelizumab for the treatment of patients with relapsed or refractory classical Hodgkin lymphoma.^3, 4^ Camrelizumab is also being investigated as a treatment for other various malignancies, including gastric/gastroesophageal junction cancer, hepatocellular carcinoma, and nasopharyngeal cancer.^5-7^

The pharmacokinetics (PK) characteristics of camrelizumab are consistent with other typical IgG4 antibodies.^8^ Non-compartmental analysis indicated a half-life of 3 – 11 days from 1 mg/kg to 10 mg/kg after a single dose. While C_max_ increased proportionally with dose from 1 mg/kg to 10 mg/kg, area under the concentration-time curve (AUC) increased in a supralinear manner over the same dose range.^8^ In phase I clinical studies of 60 to 400 mg infusions of camrelizumab, the coefficient of variation of AUC was more than 30%.^9^ Therefore, it is necessary to analyze the factors that affect PK properties of camrelizumab and to investigate the effect of these factors on dosing regimen.^10^

Early clinical studies of camrelizumab employed bodyweight-based dosing strategies of 1 mg/kg to 10 mg/kg every 2 weeks (Q2W) and compared 3 mg/kg with a flat-dose regimen of 200 mg Q2W. Although the flat-dose was selected for the subsequent expansion phase based on the PK and receptor occupancy data, the relevance of body weight to the exposure of camrelizumab has not been established. A dose adjustment of camrelizumab may be required when there is a large variation in the weight of patients.^7, 11^ Population PK analysis of data obtained in patients across multiple trials was the most efficient approach to answer this question.^12^

In this study, a population-PK model of camrelizumab was developed using pooled data from four Phase I and Phase II clinical trials to evaluate the impact of covariates on exposure, to support dose recommendations in subpopulations, and to assess the adequacy of a weight-based dosing regimen.

## 2 Methods

### 2.1 Population-pharmacokinetic Data

Data from three phase I and one phase II clinical trials in patients with advanced solid tumors, melanoma, or relapsed/refractory classical Hodgkin lymphoma were pooled to conduct this population PK analysis (Table 1). A total of 133 patients were enrolled in this analysis. The three phase 1 trials (SHR-1210-101, SHR-1210-102, SHR-1210-103) and one phase 2 trial (SHR-1210-II-204) were registered at Chinese Clinical Trial Registry (CTR20160175, CTR20160207, CTR20160248, CTR20170500, respectively). All studies were carried out in accordance with principles as defined in the Declaration of Helsinki (October 2013)^13^. The protocol and all amendments were approved by the institutional review board and independent ethics committee of each trial center. Informed consent was obtained from each patient before enrollment.

**Table 1.** Summary of clinical studies used in this population-pharmacokinetic modeling study.

| Study | Dosing regimen | Indication | Number of subjects | Number of PK Samples | Scheduled PK time points |
| --- | --- | --- | --- | --- | --- |
| SHR-1210-101 | 1mg/kg, 3mg/kg, 10mg/kg and 200mg, Q2W | Advanced solid tumors | 49 | 1140 | Cycle 1: 30 min before and 0.1, 2, 6, 24, 48, 168, 336, 504 h after end of infusion on day 1<br>Cycle 2 and subsequent cycles: 30 min before and 0.1 h after end of infusion on day 1 and 15. |
| SHR-1210-102 | 60mg, 200mg and 400mg, Q2W | Advanced Melanoma | 36 | 986 | Same as above |
| SHR-1210-103 | 60mg, 200mg and 400mg, Q2W | Advanced solid tumors | 36 | 1052 | Same as above |
| SHR-1210-II-204 | 200mg, Q2W | Relapsed or refractory classical Hodgkin lymphoma | 12 | 120 | Cycle 1: 30 min before and 0.1, 2 h after end of infusion<br>Cycle 2, Cycle 4 and Cycle 6: 30 min before and 0.1 h after end of infusion |

For the assessment of camrelizumab PK, serum samples were collected at prespecified time points in each of the studies. An intensive sampling strategy was employed in the first cycle of the three phase 1 trials (SHR-1210-101, SHR-1210-102, SHR-1210-103). Subsequent cycles of the phase 1 trials and all cycles of the phase 2 trial (SHR-1210-204) employed a sparse sampling strategy. The details of the study design in each trial are listed in Table 1.

Camrelizumab concentrations were measured by enzyme linked immunosorbent assay using a calibration range of 157 - 10,000 ng/mL for the three phase 1 trials, and 180 - 10,000 ng/mL for the phase 2 trial.^7, 9^

Patients were defined as evaluable for PK analysis if they had ≥1 adequately dose and ≥1 corresponding concentration sample. Covariates with data missing for >10% of the patients were not included in the analysis. The data of covariates with data missing for ≤10% of the patients were imputed to the population median for continuous covariates and values with higher frequency for categorical covariates.

### 2.2 Population PK Analyses

Population PK models were developed using a nonlinear mixed effect modeling (NONMEM) approach, as implemented in the NONMEM software (version 7.4.0, ICON Development Solutions, Ellicott City, MD, USA) using first-order conditional estimation with interaction. Graphical and statistical analyses, including evaluation of NONMEM outputs, were performed with Perl speaks NONMEM (PsN, version 4.7.0, Department of Pharmaceutical Biosciences, Uppsala University, Sweden), R (version 3.4.1, R Foundation for Statistical Computing, Vienna, Austria), R packages Xpose (version 4.5.3, Department of Pharmaceutical Biosciences, Uppsala University, Sweden), and Pirana (version 2.9.7, Certara, Inc. Princeton, USA).

#### 2.2.1 Base Model

In the development of the structural PK model, the concentration-time data were fitted to one- and two-compartment models with linear and nonlinear clearance (CL_nonlinear_), and the suitability of the models was assessed. Nonlinear elimination pathways were explored by incorporating CL described by Michaelis–Menten kinetics^14^ (Eq. 1):

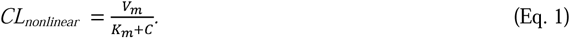

where CL_nonliear_ is the nonlinear elimination rate, V_m_ is the maximum elimination rate, C is the camrelizumab concentration, and K_m_ is the Michaelis–Menten constant, the concentration at which 50% of the maximum elimination rate is reached.

Between-subject variability (BSV) was assumed to follow a log-normal distribution and was therefore implemented into the model as follows^15^ (Eq. 2):

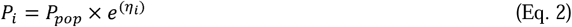

where *P*_*i*_ depicts the individual or *post hoc* value of the parameter for the *i*th individual, *P*_*pop*_ depicts the population mean for the parameter, and η_*i*_ depicts the empirical Bayes estimate of BSV for the *i*^th^ individual, sampled from a normal distribution with a mean of zero and a variance of ω^*2*^.

Residual error was evaluated as a proportional or additive error, or as a combination of both (Eq. 3).

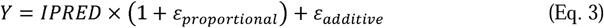

where *Y* is the observed concentration, IPRED is the individual predicted concentration, *ε*_*proportional*_ is the proportional error component, and *ε*_*additive*_ is the additive error component. Residual error components are sampled from a normal distribution with a mean of zero and variance of σ^2^.

The base model selection was based on Akaike’s information criterion (AIC)^16^, precision of parameter estimates, condition number, and goodness-of-fit plots.

#### 2.2.2 Covariate model

A three-step approach was used for the covariate analysis. In the first step, the relationship between PK parameters and covariates was screened by plotting the individual empirical Bayes estimates for PK parameters versus potential covariates. This was followed by linear regression for continuous covariates and analysis of variance testing for categorical covariates. Only those covariates with a significant effect (r>0.2, p<0.001) on the estimated PK variables, which could be meaningfully explained from both a clinical and scientific perspective, were carried through to the next stage.

In the second step, the identified covariates were added to the base model one at a time. Significance was assessed using the likelihood ratio test, where the addition of one parameter required a reduction in objective function value >3.84 (p<0.05) obtained by NONMEM during the forward inclusion. All significant covariates were included in the full model and additional covariates of borderline significance were only included if the covariate was highly likely to be influential based on scientific judgment.

The final step involved a stepwise backward elimination process, starting with the full model and removing each covariate one at a time. The covariate that was the least significant was removed and the process was repeated. The criterion for retention of a covariate in the model was a change in likelihood ratio >6.63 for one parameter (p<0.01) during the stepwise backward elimination stage.

Continuous covariates were evaluated using both a linear function and a power function (Eq. 4 and 5). Categorical covariates were tested using (Eq. 6):

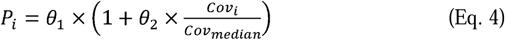

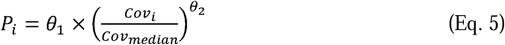

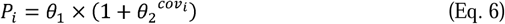

where *P*_*i*_ and *Cov*_*i*_ are the parameter and covariate value for the *i*th individual, respectively, *Cov*_*median*_ is the median value for the covariate, θ_*s*_ are the parameters to be estimated, and θ_*1*_ represents the typical value of a pharmacokinetic parameter in an individual with the median value for the covariate.

The shrinkage derived from the final model was assessed for each BSV term, as well as for residual variability.

### 2.3 Model evaluation

Goodness-of-fit plots were used for model evaluation including observed concentration (DV) vs. population predicted concentration (PRED), DV vs. individual predicted concentrations (IPRED), conditional weighted residuals (CWRES) vs. PRED, and CWRES vs. time.

The PK parameters were estimated repeatedly by fitting the final model to 1000 bootstrap datasets, sampled from the original dataset with replacement^17^. The median values and 2.5%∼97.5% of the population PK parameter estimates from these 1000 bootstrap datasets were compared with the point estimates from the final model.

The predictive performance of the final model was assessed using a visual predictive check (VPC) approach, which compared the distribution of observed concentrations and model predictions. A total of 1000 simulated datasets were generated using the final model.

### 2.4 Dosing regimen

Monte Carlo simulations, using the individual empirical Bayes PK parameters, were used to evaluate the effect of body weight on the PK of camrelizumab when administered at a dose of 200 mg Q2W and compared to the effects of a body weight-adjusted dose of 3 mg/kg Q2W.

To predict camrelizumab PK in each dosing regimen group, 1000 virtual patients were created by randomly drawing covariate values with replacement from the pooled modeling data.

BSV and residual variability in the model were sampled from the established distributions, together with PK parameters and covariate relationships for each virtual patient, which were in turn used to determine steady-state peak concentration (C_max,ss_), steady-state trough concentration (C_min,ss_), steady-state average concentration (C_average,ss_), and steady-state AUC (AUC_ss_). C_average,ss_ was calculated as (Eq. 7):

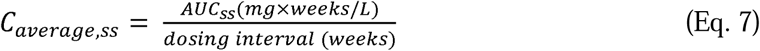

where C_max,ss_ and C_min,ss_ were determined from the concentration-time profile using each individual’s *pos-hoc* estimated pharmacokinetic parameters. AUC_ss_ is AUC in one dosing interval at steady state. Summary statistics (median, 5% – 95%) were determined using R software.

## 3 Results

### 3.1 Demographics

The dataset included 133 patients who provided a total of 3298 plasma concentrations, of which 206 samples were excluded: 203 (6.16%) samples below the limit of quantification and 3 (0.09%) samples with missed values. For covariates, no data were missing. In total, 3092 observations (93.75%) were used in the population PK analysis. A summary of patient demographics for the analysis dataset is presented in Table 2. The patients presented various tumor types, and two-third of the patients were male.

**Table 2.** Baseline demographic and disease characteristics of 133 patients.

| **Covariate** | **SHR-1210-101** | **SHR-1210-102** | **SHR-1210-103** | **SHR-1210-II-204** | **Total** |
| --- | --- | --- | --- | --- | --- |
| Number of patients | 49 (36.8%) | 36 (27.1%) | 36 (27.1%) | 12 (9.0%) | 133 (100%) |
| Number of PK Samples | 1140 (34.6%) | 986 (29.9%) | 1052 (31.9%) | 120 (3.6%) | 3298 (100%) |
| Age (years) | 47 (23 - 69) | 52 (29 - 68) | 54.5 (35 - 65) | 28.5 (21 - 50) | 50 (21 - 69) |
| Weight (kg) | 56.5 (36.8 - 72.1) | 64 (41 - 90) | 65.5 (47 - 91) | 63 (42 - 86) | 61 (36.8 - 91) |
| Sex |  |  |  |  |  |
| Male | 37 (75.5%) | 17 (47.2%) | 28 (77.8%) | 6 (50%) | 88 (66.2%) |
| Female | 12 (24.5%) | 19 (52.8%) | 8 (22.2%) | 6 (50%) | 45 (33.8%) |
| Race |  |  |  |  |  |
| Han | 49 (100%) | 34 (94.4%) | 34 (94.4%) | 11 (91.7%) | 128 (96.2%) |
| Others | 0 (100%) | 2 (5.6%) | 2 (5.6%) | 1 (8.3%) | 5 (3.8%) |
| Creatinine clearance (mL/min) | 89.07 (51.5 - 159.0) | 108.33 (52.8 - 178.7) | 101.1 (61.3 - 160.9) | 136.93 (110.7 - 210.8) | 100.69 (51.5 - 210.8) |
| Aspartate aminotransferase (U/L) | 15.4 (6.4 - 72.8) | 23.5 (13 - 82) | 21 (12 - 49) | 19 (13 - 38) | 21.7 (8 - 115.4) |
| Alanine aminotransferase (U/L) | 22.9 (8 - 115.4) | 16 (5 - 88) | 15 (7 - 55) | 13 (5 - 54) | 15 (5 - 88) |
| Total bilirubin (umol/L) | 8.3 (5.1 - 20.6) | 11.45 (5.9 - 24.1) | 9.8 (4.9 - 22.3) | 11.25 (8.4 - 24.2) | 9.7 (4.9 - 24.2) |
| Albumin (g/L) | 43.2 (29.7 - 50.4) | 45.3 (32.7 - 52.5) | 44.1 (38.2 - 50.2) | 41.7 (35.3 - 48.1) | 44 (29.7 - 52.5) |
| Tumor |  |  |  |  |  |
| Nasopharyngeal carcinoma | 31 (63.3%) | / | 3 (8.3%) | / | 34 (25.6%) |
| Lung cancer | 18 (36.7%) | / | 3 (8.3%) | / | 21 (15.8%) |
| Melanoma | / | 36 (100%) | / | / | 36 (27.1%) |
| Esophageal cancer | / | / | 14 (38.9%) | / | 14 (10.1%) |
| Gastric cancer | / | / | 5 (13.9%) | / | 5 (3.8%) |
| Classical Hodgkin lymphoma | / | / | / | 12 (100%) | 12 (9.0%) |
| Others | / | / | 11 (30.6%) | / | 11 (8.2%) |
| Co-administration |  |  |  |  |  |
| Monotherapy | 48 (98.0%) | 33 (91.7%) | 36 (100%) | 12 (100%) | 129 (97.0%) |
| Combination therapy | 1 (2%) | 3 (8.3%) | / | / | 4 (3%) |

### 3.2 Population-Pharmacokinetic Model

#### 3.2.1 Base Model

Two compartment models were found to better describe the camrelizumab phramocokinetics than the one compartment model, resulting in a decrease of >300 points in AIC. Inclusion of first order and nonlinear elimination resulted in a further decrease in AIC of 60 points compared with the linear model. The model was parameterized by clearance of linear elimination (CL_linear_), inter-compartmental clearance (Q), distribution volume of central compartment (V_1_), distribution volume of peripheral compartment (V_2_), V_m_ and K_m_. The model structure is shown in Figure 1.

**Figure 1.**
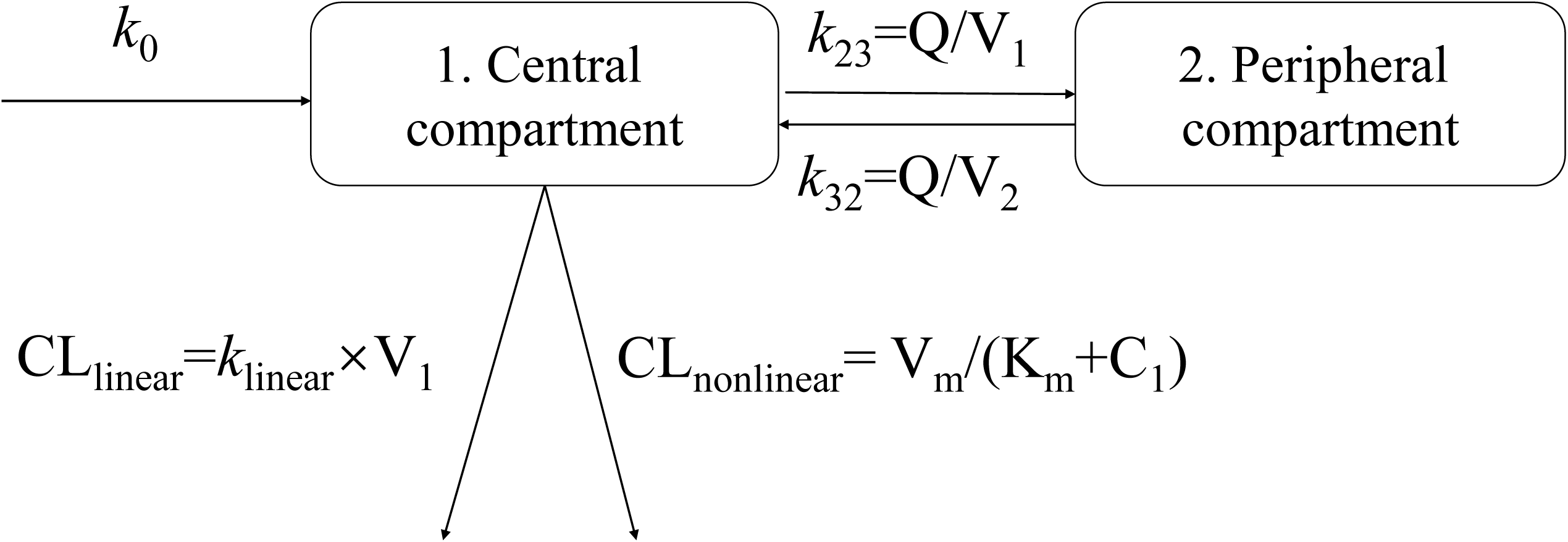
Model Structure. *k*_0_, infusion rate; *k*_23_, elimination rate from central compartment to peripheral compartment; *k*_32_, elimination rate from peripheral compartment to central compartment; *k*_linear_, linear elimination rate; CL_linear_, clearance of linear elimination; Q, inter-compartmental clearance; V_1_, apparent distribution volume of central compartment; V_2_, apparent distribution volume of peripheral compartment; *k*_nonlinear_, nonlinear elimination rate; C_1_, concentration of central compartment; V_m_, maximum elimination rate; K_m_, Michaelis–Menten constant.

BSV was estimated for CL_linear_, V_1_, and V_m_ for the acceptable precision of parameter estimates. The residual error was best described by a combined proportional and additive error model.

#### 3.2.2 Covariate Model

The covariates investigated included baseline age, weight, sex, race, creatinine clearance, aspartate aminotransferase, alanine aminotransferase, total bilirubin, albumin, hemoglobin, platelet count, and white blood cell (WBC) count. Initial graphical screening showed significant effects of albumin, hemoglobin, platelets, and WBC on CL_linear_, weight on V_1_, and weight on Q (r > 0.2, p < 0. 001). When these covariates were tested in a forward inclusion step, the effects of albumin, platelets, WBCs, and weight were significant (p < 0.05) and retained in the model. The effects of WBCs and platelets on CL_linear_, and weight on V_1_, were excluded using a stepwise backward elimination method (p > 0.01). After covariate screening, the effects of albumin on CL_linear_ and weight on Q were retained in the final model. The main steps in the covariate model building from the base model to the final model are summarized in Supplementary Materials Table S1.

The parameters of the final model are presented in Table 3. The final model is listed below (Eq. 8):

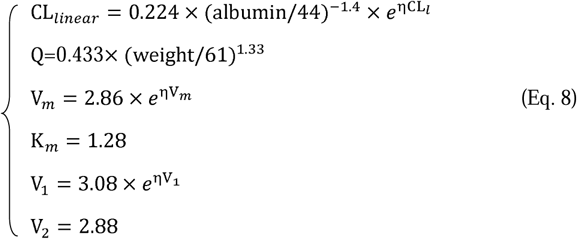

**Table 3.** Population-pharmacokinetic parameter estimates and bootstrap evaluation.

| Parameters | Base model |  | Final model |  |  |
| --- | --- | --- | --- | --- | --- |
|  | Parameter estimates<br>(%CV) | Shrinkage<br>(%) | Parameter estimates<br>(%CV) | Shrinkage<br>(%) | Bootstrap<br>Median (2.5% - 97.5%) |
| CL<sub>linear</sub> (L/day) | 0.242 (2.7) | / | 0.231 (6.1) | / | 0.23 (0.20 - 0.26) |
| V<sub>m</sub> (mg/day) | 2.86 (3) | / | 2.94 (7.5) | / | 3.00 (2.26 - 3.71) |
| K<sub>m</sub> (mg/L) | 1.28 (1.4) | / | 1.38 (13) | / | 1.40 (0.91 - 2.76) |
| V<sub>1</sub> (L) | 3.08 (2.7) | / | 3.07 (3.7) | / | 3.08 (2.77 - 3.33) |
| Q (L/day) | 0.385 (3.8) | / | 0.414 (6.7) | / | 0.41 (0.34 - 0.51) |

|  |  |  |  |  |  |
| --- | --- | --- | --- | --- | --- |
| V<sub>2</sub> (L) | 2.88 (2.8) | / | 2.9 (3.6) | / | 2.91 (2.35 - 3.35) |
| albumin on CL<sub>linear</sub> | / | / | -1.98 (24.2) | / | -1.93 (-2.94 - -0.89) |
| weight on Q | / | / | 1.22 (26.9) | / | 1.18 (0.31 - 2.28) |
| Between subject variability |  |  |  |  |  |
| CL<sub>linear</sub> (%) | 57.0 (8.6) | 11.6 | 50.8 (9) | 13 | 50.2 (32.7 - 68.9) |
| V<sub>m</sub> (%) | 48.3 (8.7) | 17.5 | 49.5 (9) | 18 | 47.8 (29.4 - 70.9) |
| V<sub>1</sub> (%) | 40.2 (6.7) | 3 | 40.7 (7) | 3 | 39.0 (17.0 - 70.68) |
| Residual variability |  |  |  |  |  |
| proportional error (%) | 29.4 (1.7) | 4.5 | 29.3 (3) | 4.5 | 28.9 (23.8 - 33.9) |
| additive error (mg/L) | 0.0812 (3.2) | 4.5 | 0.0827 (32) | 4.5 | 0.0823 (0.0293-0.112) |

CL<sub>linear</sub>, clearance of linear elimination; V<sub>m</sub>, maximum elimination rate; K<sub>m</sub>, Michaelis–Menten constant; V<sub>1</sub>, distribution volume of central compartment; Q, inter-compartmental clearance; V<sub>2</sub>, distribution volume of peripheral compartment;

CL_linear_ decreases with albumin, and a decrease from albumin 50 g/L to albumin 25 g/L is associated with a 10.3% decrease in CL_linear_; BSV was reduced from 57.0 to 50.8% for CL_linear_, indicating 10.8% of the BSV in CL_linear_ was explained by albumin; Q exhibits a linear correlation with weight.

The shrinkages of both BSV and residual variability were less than 30%, which indicates a reliable estimate of the individual empirical Bayes PK parameter (Table 3).

### 3.3 Model Evaluation

The goodness-of-fit for the final model (Figure. 2) showed a good agreement between observed and predicted values. The scatterplots of DV vs. PRED and DV vs IPRED showed random scatter around the identity line, indicating the absence of systematic bias.

**Figure 2.**
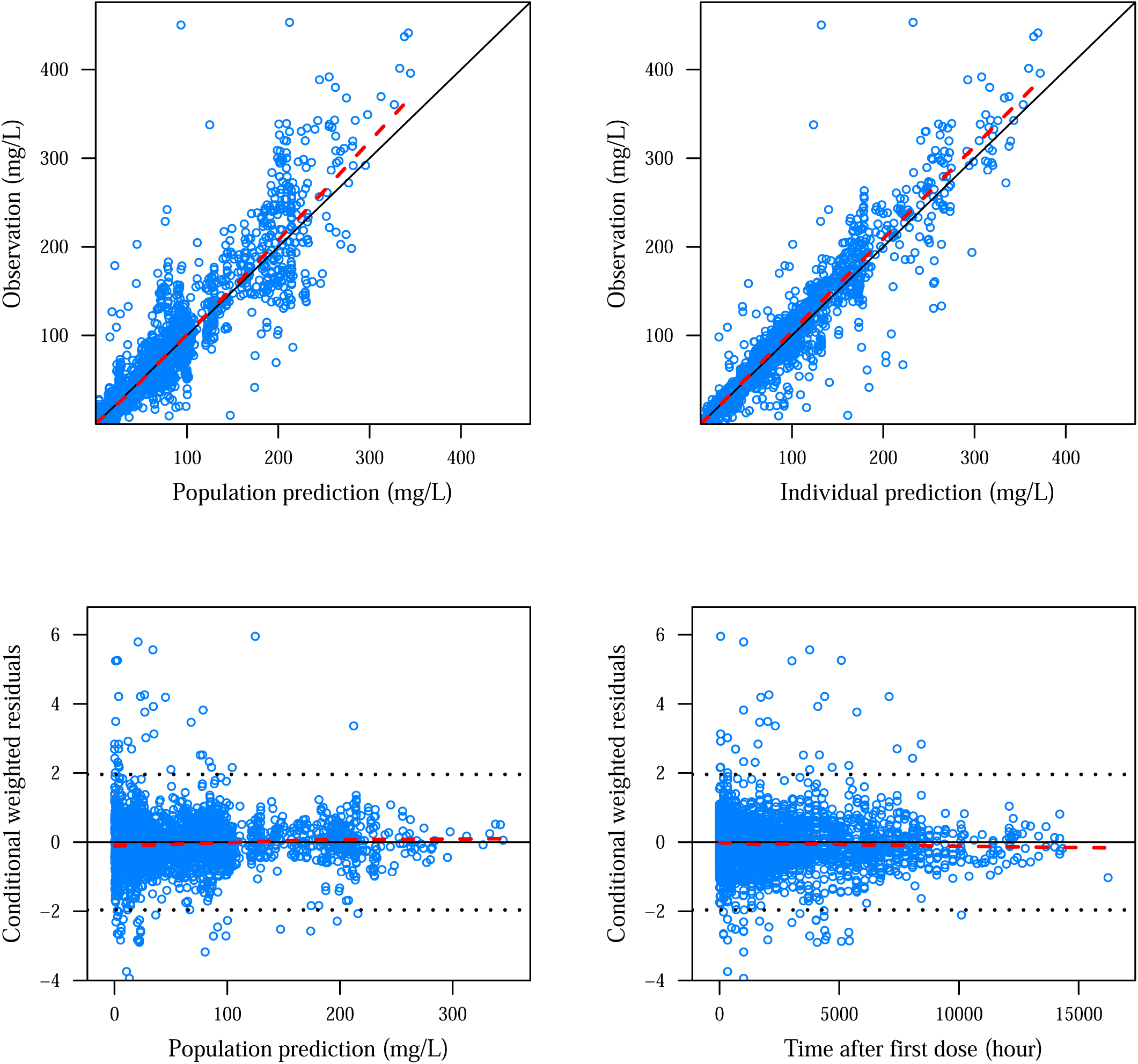
Goodness-of-fit plots of the final population-pharmacokinetic model. The red line represents the locally weighted scatterplot smoothing line.

A non-parametric bootstrap with 1000 replicates was performed for the final model, with 915 of the replicates successfully presenting the minimization step. The final model parameters and bootstrap results are presented in Table 3. Overall, the population estimates for the final model were close to the median of the bootstrap replicates and were within the 2.5 – 97.5 percentiles obtained from the bootstrap analysis. The precision of these parameter estimates was also satisfactory. The 95% CIs did not contain any null values for any parameters.

The VPC showed that the median and 95% CI of the observed data were in line with those from the simulation-based predictions from the model for all strata (Figure. 3).

**Figure 3.**
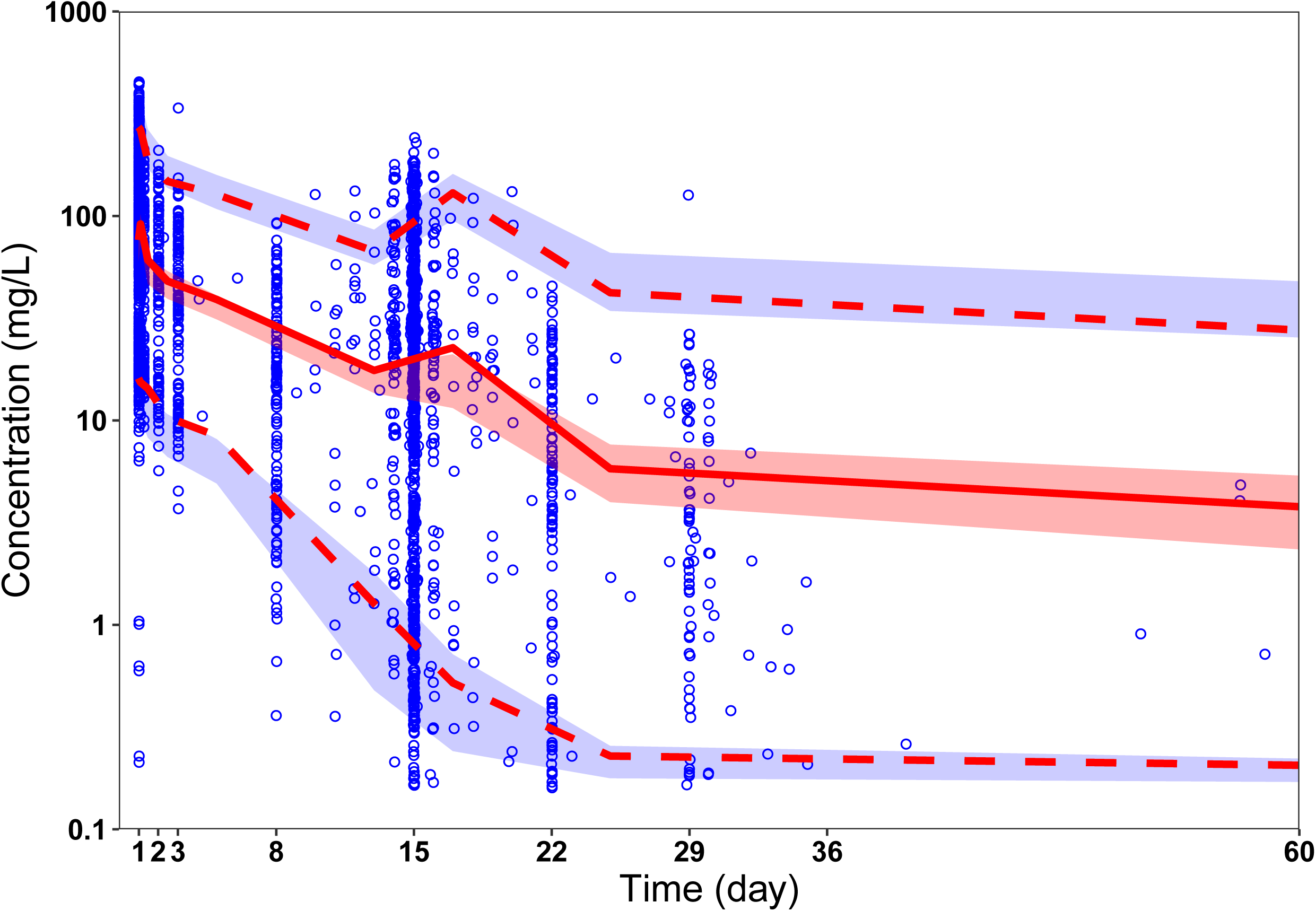
Visual predictive check. Circles represent observed data. Lines represent the 5% (dashed), 50% (solid), and 95% (dashed) percentiles of the observed data. Shaded areas represent nonparametric 95% confidence intervals about the 5% (light blue), 50% (light red), and 95% (light blue) percentiles for the corresponding model-predicted percentiles.

### 3.4 Simulations for Dosing Regimens

Summary statistics for the observed camrelizumab exposures across the 200 mg and 3 mg/kg Q2W are presented in Table 4. The 2.5% – 97.5% of C_average,ss_ for 3 mg/kg Q2W are from 12.81 to 113.87 μg/mL, which are similar to 200 mg Q2W (15.28-112.08 μg/mL). The median C_max,ss_, C_min,ss_, and C_average,ss_ values for 200 mg Q2W are higher than those for 3 mg/kg Q2W.

**Table 4.** Predicted summary statistics of camrelizumab exposure metrics.

|  | 3mg/kg every 2 weeks |  | 200mg every 2 weeks |  |
| --- | --- | --- | --- | --- |
|  | Median | 2.5%-97.5% | Median | 2.5%-97.5% |
| $C_{\max, ss}$ (µg/mL) | 89.55 | 39.27-195.41 | 96.40 | 47.26-190.36 |
| $C_{\min, ss}$ (µg/mL) | 23.11 | 1.22-92.70 | 26.13 | 1.78-90.96 |
| $C_{\text{average}, ss}$ (µg/mL)* | 41.27 | 12.81-113.87 | 45.48 | 15.28-112.08 |

$C_{\max, ss}$ , steady-state peak concentration; $C_{\min, ss}$ , steady-state trough concentration; $C_{\text{average}, ss}$ , steady-state average concentration.

## 4 Discussion

This is the first study to report a population PK model of camrelizumab in subjects with advanced melanoma, advanced solid tumors, and relapsed or refractory classical Hodgkin lymphoma. Camrelizumab PK were well described using a two-compartment model with parallel first-order and Michaelis–Menten CL from the central compartment.

The final model is in line with known characteristics of antibody PK, where the nonlinear pathway is thought to be related to clearance of the mAb via saturable target-mediated mechanisms (such as receptor-mediated endocytosis), while the linear component represents clearance pathways that are not saturable at therapeutic mAb concentrations (such as Fc-mediated elimination)^18^. The Michaelis–Menten constant in the model is 1.38 μg/mL, indicating that at low camrelizumab concentrations (<1.38 μg/mL), target-mediated elimination contributes a significant portion of the total CL. With increasing camrelizumab concentrations, the CL decreases dramatically as the target-mediated elimination pathway becomes saturated. When above the median of simulated concentration of 200 mg Q2W, the CL approaches that of the first-order process, and the contribution from the nonlinear pathway becomes negligible.

Our study also showed that camrelizumab CL decreased with increasing albumin level. The impact of albumin on PK of mAbs has been previously reported for infliximab, bevacizumab, ustekinumab, and pertuzumab.^19^ Because albumin and IgG share the same Fc receptor salvaging pathway, Fc receptor also binds and protects albumin from intracellular catabolism, thereby playing an important role in the homeostasis of both IgG and albumin.^10^ A higher albumin concentration could be an indicator of an increased number of Fc receptor s and a related reduction in the rate of camrelizumab elimination.^20^ Although albumin had a statistically significant impact on CL_linear_, simulation analyses demonstrated that the magnitude of its effect on camrelizumab exposure was limited (Figure 4). As albumin levels increased from 20 to 50 g/L, the median C_min,ss_ increased from 61 to 78 μg/mL, which are comparable to the 10 – 90% percentiles of the C_min,ss_ (39 to 123 μg/mL). Therefore, a dose adjustment for albumin is not warranted.

**Figure 4.**
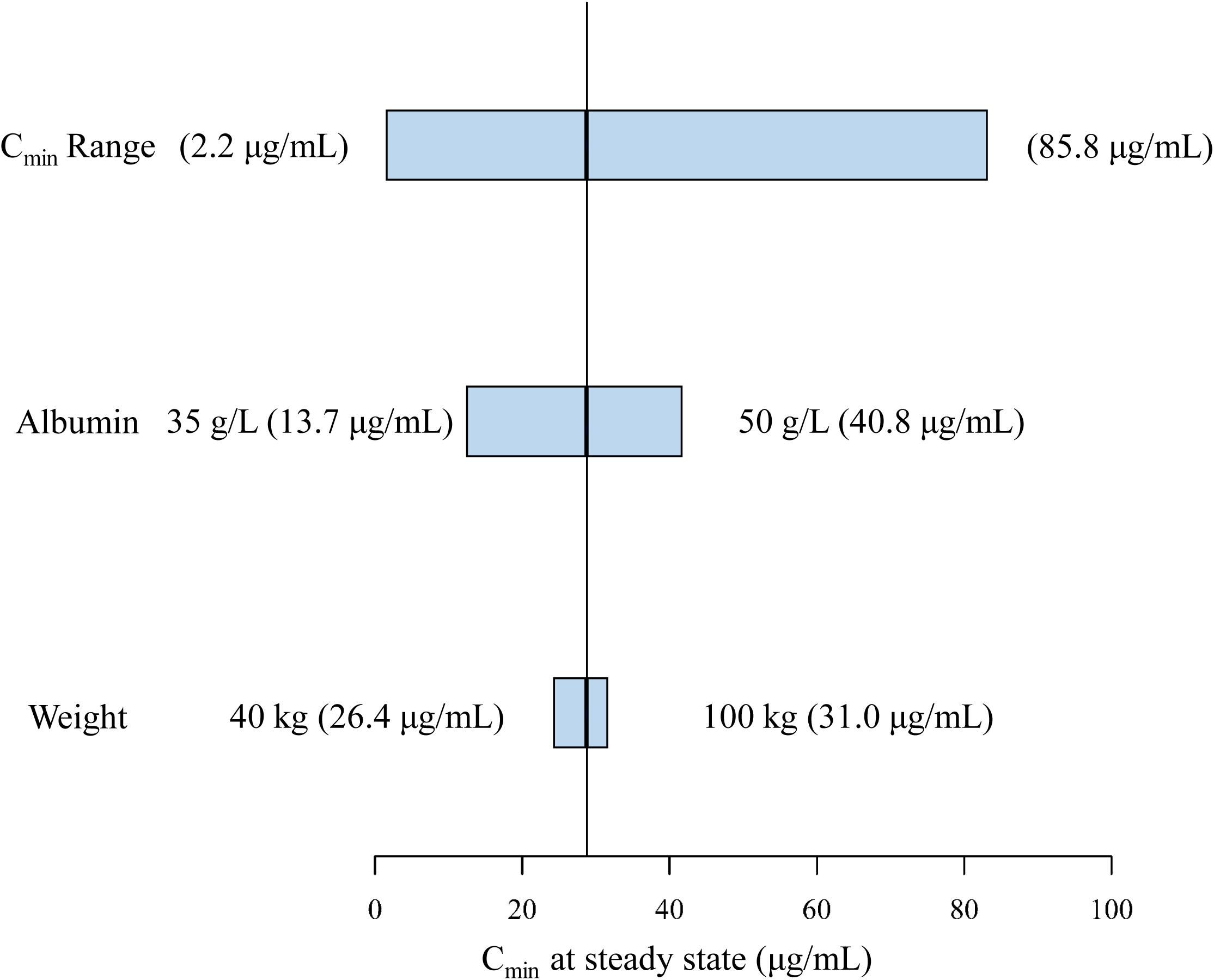
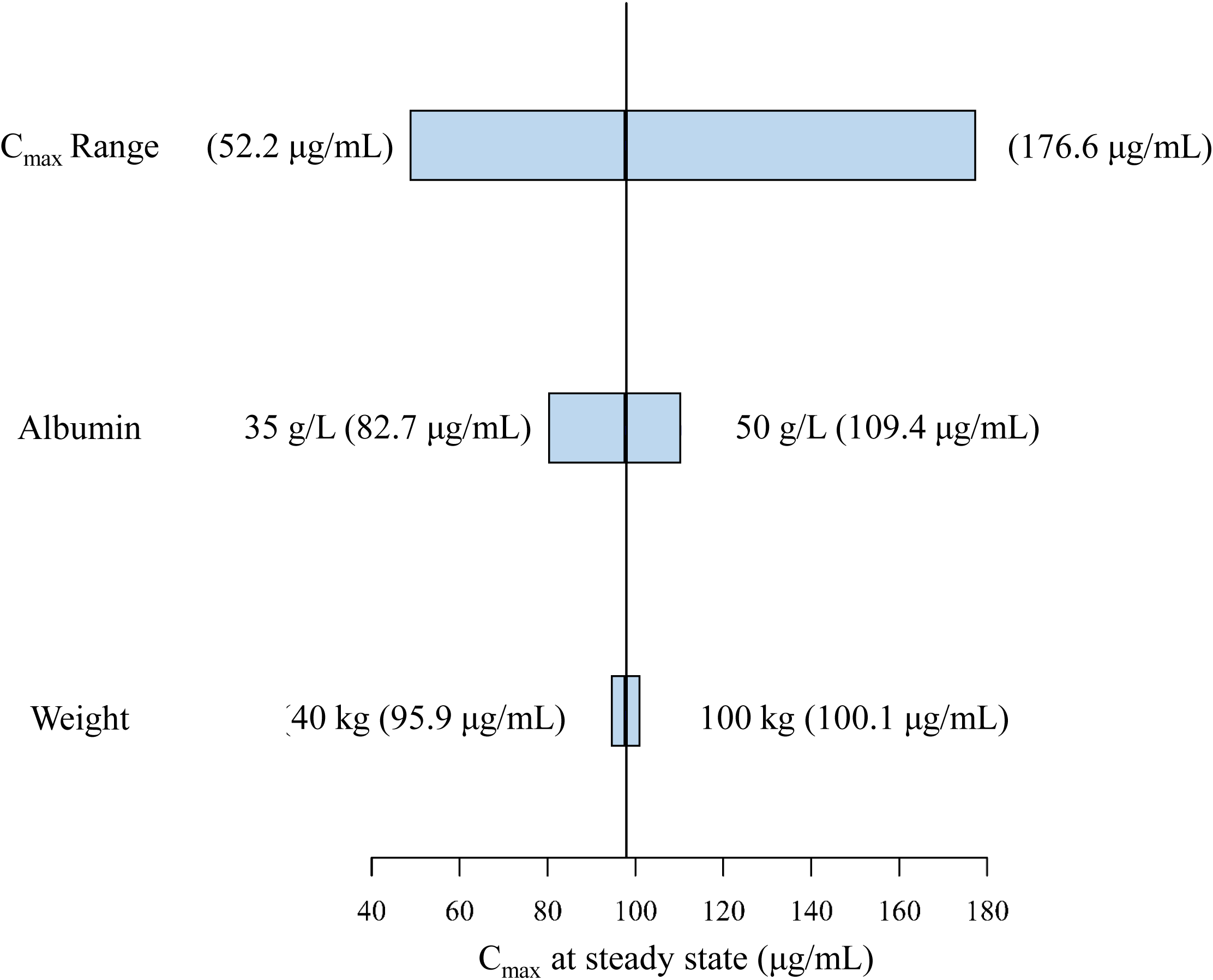
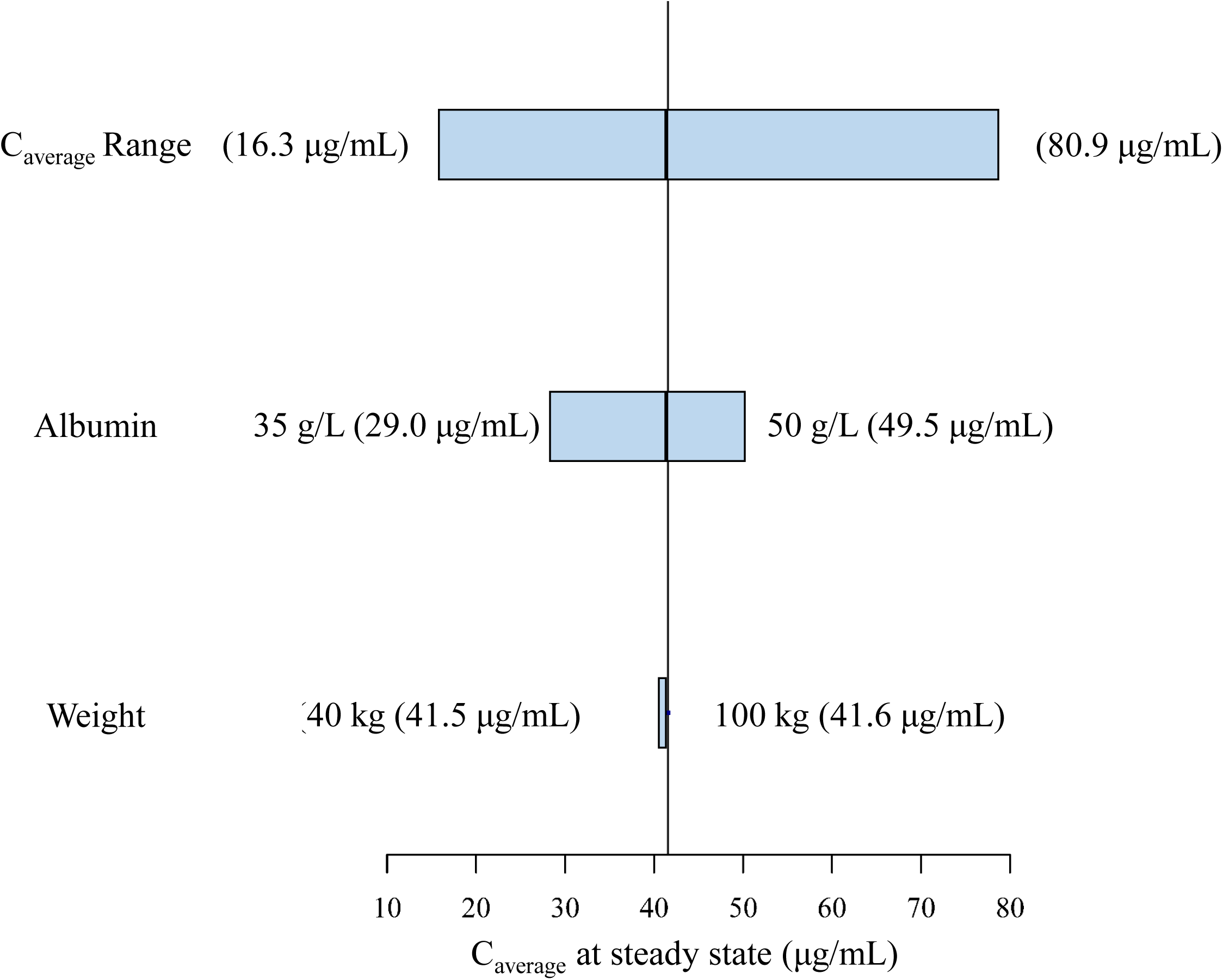
Sensitivity plots comparing effect of covariates on steady state exposure. (a) C_min_; (b) C_max_; (c) C_average_. Vertical reference lines represent typical steady-state exposure value of a 62-kg patient with albumin of 44 g/L receiving 200 mg of camrelizumab every 2 weeks. The top bars in each plot represent the 5% – 95% exposure values across the entire population. The labels at each of the lower bars indicate range of the covariate values. The length of each bar describes the impact of that particular covariate on the observed PK parameter.

Therapeutic mAb dosing is usually based on body weight^21^. However, this dosing paradigm has recently been re-evaluated because of the wide dose range for the therapeutic efficacy and tolerability for camrelizumab. Flat dose is considered and applied in the clinical settings due to increased convenience, elimination of wastage, improved safety, and improved compliance.^9^Our study showed that only weight has an impact on Q of camrelizumab, and it has little effect on camrelizumab exposure. Meanwhile, the mean exposure profile for the 200 mg flat dose is essentially similar to that of the 3 mg/kg profile. Although patients with increased weight had lower exposures with the 200 mg flat dose compared to the 3 mg/kg regimen, the distribution of exposures obtained in these patients was within the range of exposures from the prior clinical reports.^22^ It was demonstrated in this study that both weight-based and fixed flat dosing are appropriate for camrelizumab, with neither regimen providing a PK advantage over the other.

There are several limitations in this study. The study was based solely on dose-exposure analysis and all patients were from China. Whether the results could be applied to population of North American and European countries, remains to be elucidated. In addition, comprehensive safety and efficacy data were lacking. Further research including an exposure-response study is needed to inform clinical dosing strategy.

## 5 Conclusions

In this study, a population PK model of camrelizumab was developed to quantify the impact of patient characteristics on PK. The PK of camrelizumab were described by a two compartment model with parallel linear and nonlinear clearance from the central compartment. Although albumin levels and patient weight had statistically significant impacts on the PK of camrelizumab, the magnitude was limited and dose adjustments were not required. Doses of 200 mg and 3 mg/kg provided similar exposure distributions with no advantage to either dosing approach with respect to controlling PK variability.

## Abbreviations

AIC: Akaike’s information criterion;
AUC: area under the concentration-time curve;
AUC_ss_: steady-state area under the concentration-time curve;
BSV: Between-subject variability;
C: camrelizumab concentration;
C_1_: concentration of central compartment;
C_average,ss_: steady-state average concentration;
CL: clearance;
CL_linear_: clearance of linear elimination;
C_max,ss_: steady-state peak concentration;
CL_nonlinear_: clearance of nonlinear elimination;
C_min,ss_: steady-state trough concentration;
CWRES: conditional weighted residuals;
DV: observed concentration;
IPRED: individual predicted concentrations;
*k*_0_: infusion rate;
*k*_23_: elimination rate from central compartment to peripheral compartment;
*k*_32_: elimination rate from peripheral compartment to central compartment;
*k*_linear_: linear elimination rate;
*k*_nonlinear_: nonlinear elimination rate;
K_m_: Michaelis–Menten constant;
mAb: monoclonal antibody;
NONMEM: nonlinear mixed effect modeling;
PD-1: programmed cell death 1 receptor;
PK: pharmacokinetics;
PRED: population predicted concentration;
Q: inter-compartmental clearance;
Q2W: every 2 weeks;
V_m_: maximum elimination rate;
VPC: visual predictive check;
V_1_: distribution volume of central compartment;
V_2_: distribution volume of peripheral compartment;
WBC: white blood cell.

## Acknowledgements

We thank Wei-wei Wang, Qing Yang and Xiao-jing Zhang from Jiangsu Hengrui Medicine Co. Ltd for critical review of the manuscript. We would like to thank Editage (www.editage.cn) for English language editing.

## CRediT author statement

**Chen-yu Wang**: Methodology, Software, Formal analysis, Validation, Visualization, Writing – Original Draft, Writing - Review & Editing. **Chang-cheng Sheng**: Methodology, Software, Formal analysis, Writing - Review & Editing. **Guang-li Ma**: Methodology, Data Curation, Supervision, Writing - Review & Editing. **Da Xu**: Data Curation, Writing - Review & Editing. **Xiao-qin Liu**: Software, Formal analysis. **Yu-ya Wang**: Data Curation. **Li Zhang**: Data Curation. **Chuan-liang Cui**: Data Curation. **Bing-he Xu**: Data Curation. **Yu-qin Song**: Data Curation. **Jun Zhu**: Data Curation. **Zheng Jiao**: Conceptualization, Methodology, Writing – Original Draft, Writing - Review & Editing.

