## supplementary table1 for "Population Pharmacokinetics of an Anti-PD-1 Antibody Camrelizumab in Patients with Multiple tumor types and model informed dosing strategy"

**Population Pharmacokinetics of an Anti‑PD‑1 Antibody, Camrelizumab (SHR-1210), in Patients with various Tumors**

**Supplementary**

Table S1. Summary of covariate model building steps from base to final population-pharmacokinetic model of camrelizumab.
 (a) Forward Selection

| Model No. | Model description | OFV | △OFV | Reference model | Acceptance/  Significance |
| --- | --- | --- | --- | --- | --- |
| 1 | Base model | 17456 |  |  |  |
| 2 | Effect of WT on CL | 17456 | 0 | 1 | No |
| 3 | Effect of LWT on CL | 17456 | 0 | 1 | No |
| 4 | Effect of HGB on CL | 17382 | -74 | 1 | Yes |
| 5 | Effect of PLT on CL | 17392 | -64 | 1 | Yes |
| 6 | Effect of WBC on CL | 17401 | -55 | 1 | Yes |
| **7** | **Effect of ALB on CL** | **17354** | **-102** | **1** | **Yes** |
| 8 | Effect of INR on CL | 17344 | -13 | 1 | Yes |
| 9 | Effect of HGB on CL to model #7 | 17348 | -5 | 7 | Yes |
| **10** | **Effect of PLT on CL to model #7** | **17336** | **-17** | **7** | **Yes** |
| 11 | Effect of WBC on CL to model #7 | 17338 | -15 | 7 | Yes |
| 12 | Effect of INR on CL to model #7 | 17350 | -4 | 7 | Yes |
| 13 | Effect of WT on CL to model #7 | 17350 | -4 | 7 | Yes |
| 14 | Effect of LWT on CL to model #7 | 17340 | -13 | 7 | Yes |
| 15 | Effect of HGB on CL to model #10 | 17332 | -4 | 10 | Yes |
| 16 | Effect of WBC on CL to model #10 | 17331 | -5 | 10 | Yes |
| **17** | **Effect of LWT on Vc to model #10** | **17323** | **-13** | **10** | **Yes** |
| 18 | Effect of APTT on V to model #10 | 17331 | -5 | 10 | Yes |
| 19 | Effect of ADA on Vm to model #10 | 17331 | -5 | 10 | Yes |
| 20 | Effect of WBC on CL to model #17 | 17317 | -6 | 17 | Yes |
| **21** | **Effect of APTT on V to model #17** | **17316** | **-7** | **17** | **Yes** |
| 22 | Effect of ADA on Vm to model #17 | 17318 | -5 | 17 | Yes |
| **23** | **Effect of WBC on CL to model #21** | **17311** | **-5** | **21** | **Yes** |
| 24 | Effect of ADA on Vm to model #21 | 17311 | -5 | 21 | Yes |
| **25** | **Effect of ADA on Vm to model #23** | **17306** | **-5** | **23** | **Yes** |
| **26** | **Effect of WT on Q to model #25** | **17289** | **-17** | **25** | **Yes** |

(b) Backward Selection

| Model No. | Model description | OFV | △OFV | Reference model | Acceptance/  Significance |
| --- | --- | --- | --- | --- | --- |
| 27 | Remove effect of ALB on CL from model #26 | 17340 | 51 | 26 | No |
| 28 | Remove effect of PLT on CL from model #26 | 17298 | 8 | 26 | No |
| 29 | Remove effect of LWT on Vc from model #26 | 17303 | 14 | 26 | No |
| 30 | Remove effect of APTT on V from model #26 | 17296 | 6 | 26 | Yes |
| 31 | Remove effect of WBC on CL from model #26 | 17293 | 3 | 26 | Yes |
| **32** | **Remove effect of ADA on Vm from model #26** | **17293** | **3** | **26** | **Yes** |
| 33 | Remove effect of WT on Q from model #26 | 17306 | 17 | 26 | Yes |
| 34 | Remove effect of ALB on CL from model #32 | 17345 | 52 | 32 | No |
| 35 | Remove effect of PLT on CL from model #32 | 17302 | 9 | 32 | No |
| 36 | Remove effect of LWT on Vc from model #32 | 17307 | 14 | 32 | No |
| 37 | Remove effect of APTT on V from model #32 | 17300 | 7 | 32 | No |
| **38** | **Remove effect of WBC on CL from model #32** | **17297** | **4** | **32** | **Yes** |
| 39 | Remove effect of WT on Q from model #32 | 17311 | 17 | 32 | No |
| 40 | Remove effect of ALB on CL from model #38 | 17356 | 59 | 38 | No |
| 41 | Remove effect of PLT on CL from model #38 | 17315 | 17 | 38 | No |
| 42 | Remove effect of LWT on Vc from model #38 | 17312 | 14 | 38 | No |
| **43** | **Remove effect of APTT on V from model #38** | **17304** | **6** | **38** | **Yes** |
| 44 | Remove effect of WT on Q from model #38 | 17316 | 18 | 38 | No |
| 45 | Remove effect of ALB on CL from model #43 | 17363 | 58 | 43 | No |
| 46 | Remove effect of PLT on CL from model #43 | 17322 | 18 | 43 | No |
| 47 | Remove effect of LWT on Vc from model #43 | 17316 | 12 | 43 | No |
| 48 | Remove effect of WT on Q from model #43 | 17323 | 18 | 43 | No |
